# Rapid isolation of respiring skeletal muscle mitochondria using nitrogen cavitation

**DOI:** 10.1101/2022.10.05.510939

**Authors:** Awais Z. Younis, Gareth G. Lavery, Mark Christian, Craig L. Doig

**Affiliations:** School of Science and Technology, Department of Biosciences, Nottingham Trent University, Nottingham, NG11 8NS. United Kingdom

**Keywords:** mitochondria, cavitation, skeletal muscle

## Abstract

Methods of isolating mitochondria commonly utilize mechanical force and shear stress to homogenize tissue followed by purification by multiple rounds of ultracentrifugation. Existing protocols can be time-consuming with some physically impairing integrity of the sensitive mitochondrial double membrane.

**Methods:** Here, we describe a method for the recovery of intact, respiring mitochondria from murine skeletal muscle tissue and cell lines using nitrogen cavitation in combination with differential centrifugation.

**Results:** This protocol results in high yield, pure and respiring mitochondria without the need for purification gradients or ultracentrifugation. The protocol takes under an hour and requires limited specialised equipment. Our methodology is successful in extracting mitochondria of both cell extracts and skeletal muscle tissue. This represents an improved yield in comparison to many of the existing methods. Western blotting and electron microscopy demonstrate an enrichment of mitochondria with their ultrastructure well-preserved and an absence of contamination from cytoplasmic or nuclear fractions. Using respirometry analysis we show that mitochondria extracted from the murine skeletal muscle cell lines and tibialis anterior have an appropriate respiratory control ratio. These measures are indicative of healthy coupled mitochondria.

**Conclusion:** Our method successfully demonstrates the rapid isolation of functional mitochondria and will benefit researchers studying mitochondrial bioenergetics as well as providing greater throughput and application for time-sensitive assays.

## Introduction

Required for locomotion, respiration, and storage of nutrients, skeletal muscle depots are densely packed with mitochondria. As such, optimal mitochondrial function is fundamental to both cellular behaviour and preservation of whole-body energy homeostasis. Click or tap here to enter text.

As appreciation of tissue-specific mitochondrial biology advances, there is a growing need to develop alternative methodological approaches to rapidly isolate and evaluate mitochondrial phenotype. However, to examine respiratory function isolation from whole-cell constituents must be both efficient and robust. Currently, this can be achieved using a variety of techniques such as differential centrifugation (5), affinity purification (6)and free-flow electrophoresis/field-flow fractionation (Islinger et al., 2010). However, there are limitations to these methods including reduced purity and yield. Moreover, certain procedures require costly reagents (8,9). Reproducible lysis of cellular membranes between samples is also important, particularly given the resilient nature of skeletal muscle cell membranes (Franzini-Armstrong & Engel, 2012; Navarro et al., 2017) contributes to impairing effective lysis. Commonly, this is resolved through mechanical homogenisation or application of shear stress. Whilst these are effective, they can compromise organelle integrity and produce compartment leakage. This is evidenced by a recent review (10) suggesting that mitochondria isolated using homogenisation show reduced integrity. Distinct methods of mitochondrial isolation also have specific requirements of time, technical capability, and specialist equipment, variables that contribute to differences in recovered purity and yield.

An ideal method should provide time-efficient mitochondrial isolation free from major contaminants and produce reproducible Respiratory Control Ratio (RCR) values consistent with established understanding of mitochondrial function. Here we apply nitrogen cavitation to cultured skeletal muscle cells and murine tissue, a procedure in which nitrogen gas is diffused in to cells under pressure and when released causes lysis of cell membranes for subsequent mitochondrial isolation. (11). This coupled to a rapid differential centrifugation process allowed recovery of mitochondria in high yield, purity and suitable for high-resolution respiratory studies. Variation in RCR is low with ±0.2 and 0.8 SEM in isolated mitochondria from tissue and cells respectively. This protocol would benefit those studying mitochondria in skeletal muscle for high-resolution respiratory studies as well as for isolating mitochondria for proteome analysis.

## Methodology

### Cell culture and tissue

For cells, C2C12 murine myoblasts (ATCC) in a T175 flask were maintained in high glucos-Dulbecco’s Minimal Eagle’s Medium (DMEM) supplemented with 10% (v/v) foetal bovine serum, 1% L-glutamine (v/v) (Life Technologies, Paisley, UK), and 1% (v/v) penicillin-streptomycin (Life Technologies) in a humidified atmosphere with 5% CO_2_ at 37°C. Once at 80% confluence, cells were incubated in DMEM supplemented with 2% (v/v) heat-inactivated horse serum (Life Technologies) for 5 days to facilitate myocyte differentiation with media being replaced every 48 hours. Cells were used between passages 18-28. Skeletal muscle tissue (*tibialis anterior*) was freshly dissected from C57/BL6J and placed in ice cold sucrose buffer.

### Mitochondrial isolation

Cell pellets were collected by washing the monolayer twice with Dulbecco’s phosphate buffered saline without Ca^2+^ and Mg^2+^ (DPBS-) to remove traces of serum. Cells were then incubated with trypsin for approximately 1 minute at 37°C. Trypsin was quenched with 10mL of complete growth media, the suspension was then transferred to a 30ml centrifuge tube. Cells were pelleted by centrifugation at 300g for 5 minutes. The supernatant was discarded, and pellet was washed twice with PBS.

Once washed the pellet was resuspended in 10mL sucrose extraction buffer (10 mM HEPES, pH 7.5, 70 mM sucrose, 200 mM mannitol, 1 mM EGTA, 1% protease inhibitors and 1% phosphatase inhibitors).

Tissue was placed in a 1.5mL centrifuge tube in 1mL of mitochondrial isolation buffer and chopped into small (approx. 2mm) chunks. The tissue suspension was transferred into a pre-cooled Dounce homogeniser and homogenised with 10 strokes of a tight pestle. The suspension was strained sequentially through 70µm and 40µm cell strainers. The strained solution was made up to 10mL with sucrose buffer.

Both cell and tissue extracts were then subject to the same steps from this point onwards. The cell suspension was placed in a nitrogen cavitation chamber (labelled a in figure 1B) with a magnetic stirrer and was pressurised to 500psi for 10 minutes whilst stirring on ice. The suspension was released drop wise through the outlet valve (labelled b in figure 1B) into a pre-chilled 15ml tube. The cell suspension was centrifuged at 800g for 10 minutes to pellet nuclei and cellular debris. The supernatant was transferred to a pre-chilled tube and the pellet discarded. The suspension was centrifuged at 11,000g for 15 minutes. The resulting pellet was resuspended in 1mL of sucrose buffer and centrifuged again at 11,000g for 10 minutes to wash the mitochondrial pellet. The method is summarised in figure 1A.

**Figure 1.**
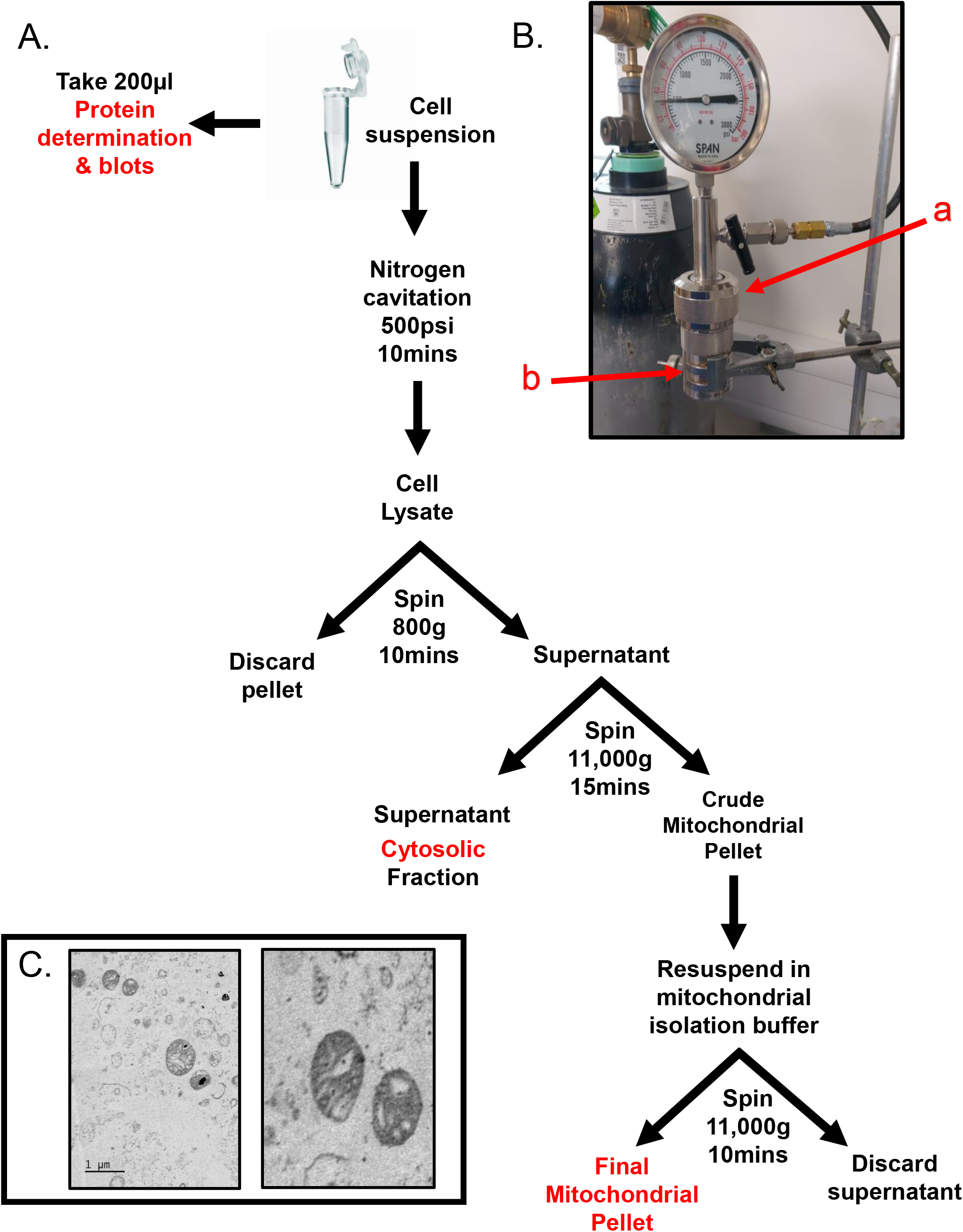
Rapid and Intact skeletal muscle mitochondrial extraction through nitrogen cavitation and differential centrifugation. **A**. Schematic of mitochondrial isolation protocol for cells or skeletal muscle tissue. B. Nitrogen cavitation device, label a (red) identifies the cavitation chamber, label b (red) identifies the outlet valve. C. Electron microscopy of recovered mitochondria at 1µm (Left) and 500nm (Right). The mitochondrial inner and outer membranes and cristae are clearly visible. The mitochondrial matrix shows no signs of damage or compromised integrity.

The pellet was then resuspended in 200µL mitochondrial assay buffer (MAS) (70mM sucrose, 210mM mannitol, 2mM HEPES, 1mM EGTA, 5mM magnesium chloride, 10mM potassium dihydrogen phosphate, 0.2% fatty acid-free bovine serum albumin (w/v) 10mM sodium pyruvate, 2mM D-Malic acid) and protein content was quantified using a modified Lowry method.

### Isolated Mitochondria Respiratory Studies

Mitochondrial respiratory studies were performed to the direction of the protocol described by (Sakamuri et al., 2018). Briefly, the SeahorseXF24 sensor cartridge was loaded with ADP 20mM (port A), oligomycin 50µM (port B), Carbonyl cyanide-p-trifluoromethoxyphenylhydrazone (FCCP) 50µM (port C) and rotenone 100µM/antimycin A 20µM (port D). Isolated mitochondria were diluted to the desired final concentration (5μg/50μL) using ice-cold MAS and loaded into the wells. The cell culture microplate was centrifuged at 2,000×g for 20 mins at 4°C. After centrifugation pre-warmed 150μL of MAS was added to each well and the plate incubated in a 37°C non-CO_2_ incubator for 10 minutes.

The plate was loaded on to the Seahorse apparatus and the injection protocol summarised in table 1 was set up.

**Table 1.**
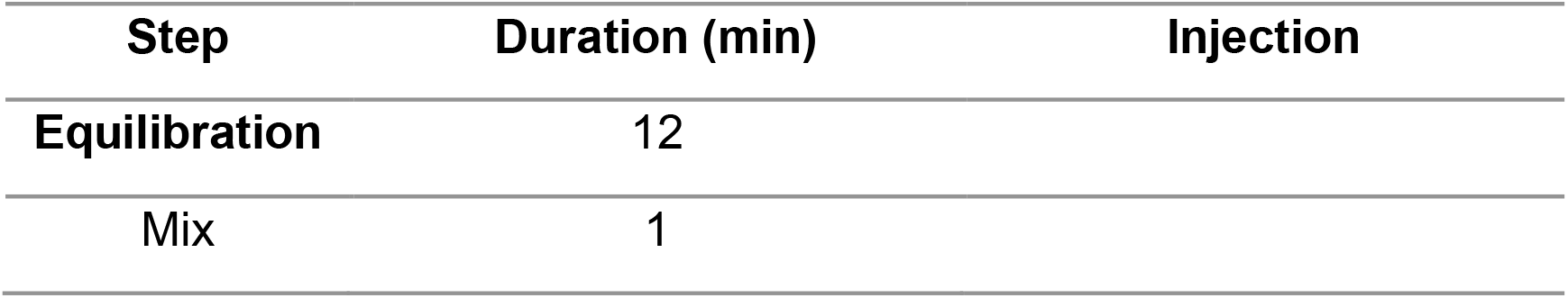
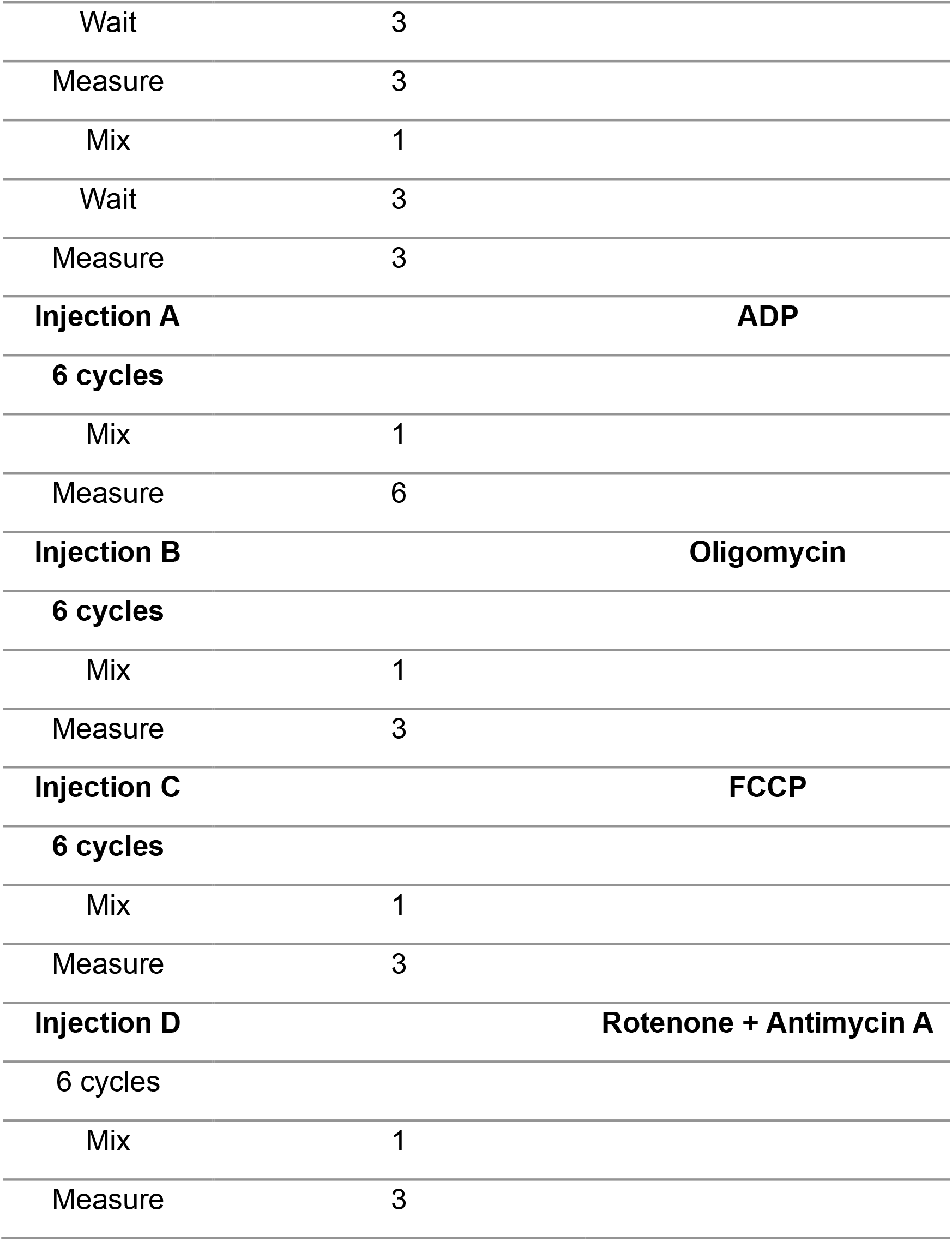
Seahorse XFe24 assay protocol for mitochondria isolated from skeletal muscle cell lines and tibialis anterior tissue.

### Western blot analysis

#### Sample preparation for Western blotting

Cells were detached using trypsin, centrifuged at 1200 rpm for 5 minutes and washed with Dulbecco’s modified Eagle’s medium. Cell pellets were weighed and resuspended in 8X pellet weight of ice-cold RIPA buffer (150mM NaCl, 5mM EDTA, 50mM Tris pH 8.0, 1% NP40, 0.5% sodium deoxycholate, 0.1% sodium dodecyl sulfate). Lysates were incubated on ice for 15 minutes with regular vortex mixing. Protein lysates were then centrifuged at 14,000g for 10 minutes at 4°C and supernatant collected. Protein content of samples was quantified using the Bio-Rad DC protein assay kit following the manufacturer’s instructions. Organelle extracts were kept in their original extraction buffer and were subject to protein content analysis as described.

#### Western blotting

Equal protein aliquots per sample were subjected to electrophoresis on either a 12% or 10% sodium dodecyl sulfate-polyacrylamide gel. Separated proteins were transferred onto a nitrocellulose membrane using the Bio-Rad Trans-Blot Turbo semi-dry blotting system. Equal protein loading was assessed by staining with 0.05% copper phthalocyanine in 12 mM HCl. Blotted membranes were blocked for 1 hour in 3% dried skimmed milk in Tris-buffered saline (TBS) containing 0.1% Tween-20 and incubated overnight at 4°C with primary antibodies. Membranes were then washed and incubated for 2 hours at room temperature with horseradish peroxidise-conjugated anti-rabbit/mouse immunoglobulin G secondary antibodies. Antibody binding was revealed with enhanced chemiluminescence western blotting detection reagent (ThermoFisher). Digital images were captured using a SYNGENE GBOX and band intensity was quantified using image J with band intensity normalised to total protein (quantified using copper phthalocyanine).

### Transmission Electron Microscopy

Samples were prepared according to the general procedure as described by (12) but with immediate processing without storage. The resin was TAAB 813 (TAAB, Aldermaston) and after sectioning samples were stained with EM Stain 336 (Agar Scientific Ltd., Stansted) and Reynold’s stain. Once fixed, embedded, sectioned, and stained, sections were examined with a JEM2100Plus (JEOL UK, Welwyn Garden City) operating at 120 kV and operated according to the manufacturer’s procedures. Electron micrographs were digitised using a Rio16 (Gatan UK, Abingdon) camera operated using Digital Micrograph (3.32.2403.0) and exported to .tiff for further analysis.

## Results

The isolation of mitochondria through nitrogen cavitation takes approximately 45 minutes. Total protein yield from a 330mg of cell pellet was 830µg of mitochondrial protein (Table 2). This equates to 0.28% of the input protein. As mitochondria are reported to make up between 3-5% of skeletal muscle, our method is successful in isolating nearly 30% of the cell’s mitochondria (13). From tissue, the method was successful in isolating 358µg of mitochondria from an average tissue weight of 107mg representing 0.34% of mitochondrial recovery from total tissue input (Table 2).

**Table 2.** Mitochondrial protein yields.

|  | **Total Starting Pellet Weight (mg)** | **Mitochondrial Protein Yield (µg)** |
| --- | --- | --- |
| **Cells** | 327.27 ±13.21 | 847.97 ± 52.27 |
| **Tissue** | 107.67 ± 6.69 | 357.79 ± 29.66 |

Isolated mitochondria from both tissue and cells showed no contamination from cytosolic or nuclear proteins showing a pure fraction without contamination from other major organelles. Western blot analysis of cytochrome C and COX IV showed enrichment in the mitochondrial fraction (figure 2A+B). Analysis of cytochrome C indicates that the mitochondria were intact with an absence of cytochrome C leakage into the cytosol. (figure 2A+B).

**Figure 2.**
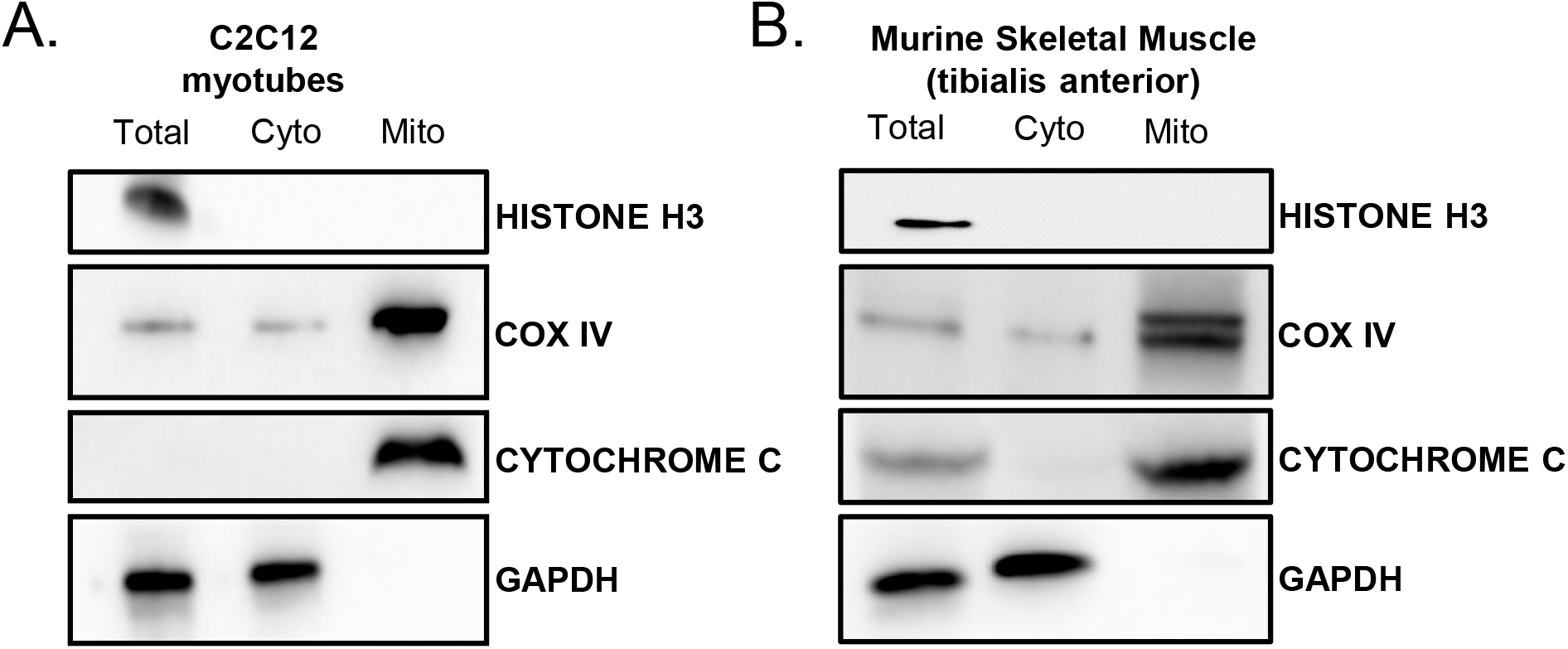
Nitrogen cavitation with differential centrifugation is successful in isolating intact, contaminant free, respiring mitochondria. A-B. Western blot analysis of nuclear (Histone H3), cytosolic (GAPDH) and mitochondrial fractions (cytochrome C and COX IV) from c2c12 cells and B tibialis anterior. C-D. Electron microscopy of isolated mitochondria. All blots have been cropped to minimise white space.

States of respiration in the isolated mitochondria from both tissue and cells were measured using a Seahorse XFe24 analyser in the presence of complex I substrates malate and pyruvate. Isolated mitochondria responded to complex I, III and V inhibitors (figure 3A+B).

**Figure 3.**
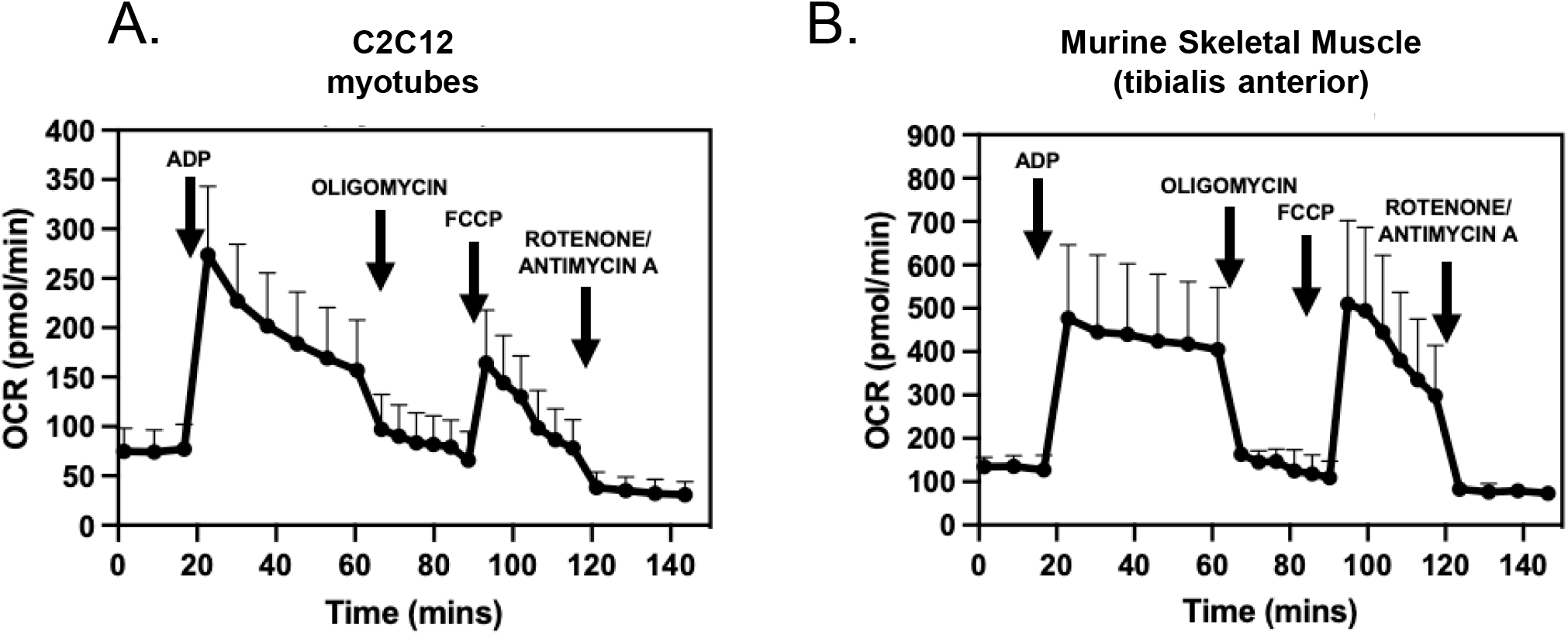
OCR curves of mitochondria isolated for A. tibialis anterior and B. C2C12 cells. Respiratory parameters of isolated mitochondria in the presence of glutamate and malate (0.5M/0.5M). The rates of respiration in State 3 and in State 4 are expressed as pmol O2/min. Data represented as a mean ± S.E.M. of six different mitochondrial preparations isolated from different skeletal muscle cells and three different TA preparations. Representative plot of point-to-point OCR data in the presence of mitochondrial substrates and inhibitors. Data expressed as a mean of 5 individual replicates + SEM.

These data demonstrate the method was able to preserve mitochondrial respiratory activity. The respiratory control ratio of isolated mitochondria from C2C12 myoblast cells was 4.9 ± 0.8 in the presence of malate and glutamate and for skeletal muscle tissue was 4.6 ± 0.2. State III respiration was measured at an oxygen consumption rate (OCR) of 273.8 ± 69.20 pmol/min and state IV respiration measured at an average OCR of 71.4 pmol/min ± 26.70 for mitochondria isolated from cells. For mitochondria isolated from skeletal muscle tissue, state III respiration was measured at an OCR of 480.27 ± 174.87 pmol/min and state IV at 107.72 ± 39.78 pmol/min (table 3).

**Table 3.** Respiration states of isolated mitochondria.

|  | State III | State IV | Respiratory Control Ratio |
| --- | --- | --- | --- |
| Cells | $273.80 \pm 69.20$ | $71.40 \pm 26.70$ | $4.90 \pm 0.80$ |
| Tissue | $480.27 \pm 174.87$ | $107.72 \pm 39.78$ | $4.55 \pm 0.17$ |

## Discussion

Selection of a mitochondrial extraction protocol is done to best suit the demands of the upstream experimental workflow and downstream endpoints. Herein, we provide an additional protocol for the rapid and robust isolation of mitochondria. This would be ideal for studies of a larger scale or those requiring higher throughput respirometry analysis.

Direct comparisons between different methodologies are challenging due to inherent variation such as cell type, chemical reagents, equipment and methodological disparities. However, utilised protocols for mitochondrial isolation each have their own limitations (9,14,15).

Click or tap here to enter text.Click or tap here to enter text.Click or tap here to enter text.Click or tap here to enter text.Click or tap here to enter text.Click or tap here to enter text.Click or tap here to enter text.Click or tap here to enter text.Click or tap here to enter text.Click or tap here to enter text.Click or tap here to enter text.Our method has the benefits of a consistent cellular lysis followed by a brief differential centrifugation to isolate intact, respiring mitochondria with a comparable RCR at the top end of the range stated in literature (14,15,22,23). Click or tap here to enter text.As expected, and evident in other methods, lysosomal and endoplasmic reticulum (ER) contamination are apparent using our method. However, there are no nuclear, cytoplasmic or whole cell contaminants (Figures 2A+B). As there are several contacts sites between mitochondria and the ER it is not unexpected that there is some ER present (30–32). In this regard, our method for isolating mitochondria may hold significant advantages for mitochondrial proteomic analysis. Due to technological advances of LC-MS/MS and availability of up-to-date and in-depth databases including MitoCarta, it may be unnecessary to have ultrapure mitochondrial fractions as proteins can be matched to these databases and other proteins from contaminants, such as lysosomes, filtered out. Therefore, we present a valuable alternative to existing mitochondrial isolation protocols. This protocol is applicable to tissue and cells of skeletal muscle origin and will be of potential use to those working with *in vivo* primary endpoints that have downstream *in vitro* mitochondrial evaluation.

## List of abbreviations

OCR: Oxygen consumption rate
MAS: mitochondrial assay buffer
FCCP: Carbonyl cyanide-p-trifluoromethoxyphenylhydrazone
TBS: Tris-buffered saline
RCR: Respiratory control ratio

## Declarations

### Ethics approval and consent to participate

Not applicable

### Consent for publication

Not applicable

### Availability of data and materials

The datasets used and/or analysed during the current study are available from the corresponding author on reasonable request.

### Author contributions

Concept and experimental design: AZY, MC and CLD. Acquisition, analysis and interpretation of datasets AZY, GGL, MC and CLD. Manuscript preparation and editing AZY, GGL, MC and CLD.

## Acknowledgments

The authors would like to thank Dr Graham J. Hickman for TEM sample preparation and imaging. Electron microscopy data used in this contribution was provided by the Imaging Suite at the School of Science & Technology at Nottingham Trent University.

